# Multi‑Stage Singular Value Decomposition for Ultrafast Ultrasound Imaging of Microbubbles

**DOI:** 10.64898/2026.05.04.722634

**Authors:** Ge Zhang, Henri Leroy, Rideau Benoît, Alec Reygrobellet, Mathieu Pernot, Thomas Deffieux, Nathalie Ialy-Radio, Sophie Pezet, Mickael Tanter

## Abstract

Microbubble contrast-enhanced ultrasound (CEUS) relies on discriminating nonlinear bubble signals from linear tissue backscattering. While Singular Value Decomposition (SVD) filtering improves this discrimination, existing techniques often fail to retain the slowly-moving microbubble signals from static clutter. Here, we present a novel multi-stage singular value decomposition (MS-SVD) framework for ultrafast CEUS imaging. Our method employs plane-wave transmissions at multiple angles and acoustic pressure levels (implemented via duty-cycle modulation) and alternating transmit polarity. The beamformed data are then processed by three sequential SVD filters: (1) spatial-angular SVD to extract coherent signals across all transmit angles, (2) spatial-pressure SVD to separate linear fundamental and nonlinear harmonic components, and (3) spatiotemporal SVD to isolate moving microbubble echoes from tissue clutter. In *in vitro* flow phantoms and *in vivo* rat brain through a cranial window, MS-SVD dramatically improves microbubble detection compared to conventional SVD filtering, MS-SVD yields much stronger vascular contrast and suppresses tissue clutter to a greater extent. The resulting power-Doppler and super-resolution maps are notably cleaner and more complete: MS-SVD detects substantially more microbubble events in ULM, revealing finer vessel details and more accurate flow speeds. By capturing the full acoustic signature of microbubbles (both fundamental and harmonic), MS-SVD achieves higher contrast-to-noise and sensitivity in CEUS. These gains make it a powerful front-end for super-resolution ultrasound localization microscopy and other high-sensitivity microvascular imaging applications.

## 1. Introduction

Ultrasound imaging has emerged as an indispensable modality in biomedical applications, offering non-invasive, real-time visualization with high spatiotemporal resolution [1, 2]. The introduction of contrast-enhanced ultrasound (CEUS) with microbubble contrast agents has substantially advanced the capability to visualize microvascular architecture and assess tissue perfusion dynamics [3-5]. These microbubbles, typically comprising a gas-filled core encapsulated by a stabilizing lipid or protein shell with diameters ranging from 1 to 10 μm, exhibit characteristic nonlinear acoustic behavior when insonified with ultrasound beyond specific pressure thresholds [4]. This pressure-dependent oscillation generates robust nonlinear backscatter signals detectable by ultrasound transducers, providing microbubble-specific contrast that can be leveraged to enhance imaging sensitivity and specificity [6, 7].

Over the past decade, singular value decomposition (SVD) of raw ultrasound data has demonstrated superior performance compared to conventional filtering approaches across multiple ultrasound imaging applications. In ultrafast ultrasound, spatiotemporal SVD filtering has enabled unprecedented discrimination between tissue motion and blood flow based on differences in spatiotemporal coherence, yielding ultrasensitive Doppler imaging capabilities [8]. The spatial-angular SVD beamformer has been successfully applied to ultrafast compounded plane-wave acquisitions, facilitating adaptive aberration correction by estimating optimal phase correction profiles required to synthesize coherent images with different plane wave transmit angles [9] or different diverging waves transmissions [10]. Furthermore, SVD beamforming was applied successfully to retrieve sound speed estimates in ultrasound data and applied successfully in the framework of liver steatosis [11, 12]. SVD beamforming can also be applied to recombine ultrasound images acquired with distinct probes [13, 14]. In the field of nonlinear imaging, higher-order SVD of ultrasonic signals acquired at different pressure levels and different temporal acquisitions was recently proposed to detect moving microbubbles [15].

Building upon the foundations of spatiotemporal SVD filtering and spatial-angular SVD beamforming, we previously introduced spatial-pressure amplitude-modulated SVD (AM-SVD), an adaptive CEUS processing technique that synergistically combines SVD decomposition with plane-wave transmissions at varying acoustic pressures, modulated through duty cycle adjustments, across multiple steering angles [16]. By exploiting the differential acoustic responses of tissue, contrast agents, and noise to systematic pressure variations, AM-SVD decomposes acquired raw data into constituent singular vectors, enabling selective reconstruction of contrast-specific signals with markedly enhanced signal-to-background ratio (SBR). Despite its demonstrated efficacy, AM-SVD operates exclusively in the fundamental frequency domain, leaving unexplored its potential for second-harmonic imaging applications. To further capitalize on the harmonic signatures of contrast agents, we subsequently developed harmonic amplitude-modulated SVD (HAM-SVD), which integrates the principles of AM-SVD within the nonlinear harmonic imaging framework [17]. HAM-SVD leverages complementary advantages from both AM-SVD and nonlinear harmonic imaging techniques to effectively isolate second harmonic components through phase-modulated transmission sequences, while AM-SVD efficiently suppresses residual tissue clutter signals that pulse inversion alone cannot completely eliminate. This dual capability is achieved by transmitting multiple amplitude-modulated pulses with alternating polarity. Preliminary investigations demonstrate that HAM-SVD provides superior tissue background suppression and enhanced isolation of second-harmonic signals from gas vesicles compared to AM-SVD, suggesting considerable promise for preclinical and translational CEUS applications.

However, several critical knowledge gaps remain. First, while previous studies of AM-SVD and HAM-SVD focused on gas vesicles - genetically encodable protein nanostructures that serve as biomolecular ultrasound reporters - their performance characteristics with larger conventional lipid-shelled microbubble contrast agents of micrometric size remain uncharacterized. Given that microbubbles exhibit substantial second harmonic generation capability, HAM-SVD is hypothesized to further optimize microbubble-specific contrast signals. Second, previous implementations of AM-SVD and HAM-SVD exploited only spatial-pressure coherence within individual image frames; the potential of spatiotemporal and spatial-pressure coherence analysis applied to time-series HAM-SVD datasets has not been explored. Third, a persistent challenge in SVD-based spatiotemporal filtering is the inadvertent suppression of slowly moving microbubbles due to increased spatiotemporal coherence with tissue clutter [18]. The synergistic combination of spatial-angular, spatial-pressure and spatiotemporal filtering strategies for flowing microbubble populations warrants systematic investigation.

Here, we addressed these gaps by introducing a multi-stage, multi-domain SVD processing pipeline for ultrafast microbubble imaging. We acquired data using coherent plane-wave compounding (5 angles per transmit burst) at four duty cycles and two polarities, yielding a dataset rich in angular and pressure diversity. In post-processing, we apply three cascaded SVD steps:

1. ***Spatial-angular SVD:*** We reshaped the data to combine all angles and extract the most coherent angular component. This step both performs adaptive aberration correction and consolidates the flow information across views.
2. ***Spatial-***pressure ***SVD*** (HAM-SVD): The angle-compounded frames at different amplitudes and polarities were stacked and decomposed. This isolates distinct acoustic responses: for example, we find that one singular vector predominantly contains the linear (fundamental) component, while another captures the nonlinear (harmonic) component. This separation goes beyond simple pulse inversion: it adaptively filters residual tissue echoes that would otherwise leak into the harmonic signal.
3. ***Spatiotemporal SVD***: Finally, each separated component was temporally filtered to distinguish moving bubbles from static tissue. By applying SVD in the slow-time domain with an aggressive cut-off (keeping the few largest singular values), we further suppress background clutter while preserving even slowly moving microbubbles.

This multi-dimensional SVD approach enabled full access to both fundamental and harmonic microbubble signals. Together, these filters significantly enhance the contrast-to-noise ratio (CNR) of CEUS images. In following, we demonstrate that MS-SVD outperforms conventional SVD and basic pulse inversion methods in several key metrics: it yields clearer Doppler images, increased microbubble detection in ULM, and more accurate flow quantification, without compromising frame rate or field-of-view.

## 2. Materials and Methods

### 2.1. *In Vitro* Wall-Less Flow Phantom Imaging

An agarose phantom consisting of a Y-shaped junction was utilized to test the multi-stage SVD sequence. The Y-shaped junction has an inlet of a diameter of 0.8 mm and two outlets with a diameter of 0.4 mm. The linear ultrasound transducer (128 elements, 15.625 MHz central frequency and a 67% bandwidth at −6 dB) equipped with ultrasound research platform (Verasonics Vantage 256, Kirkland, WA, USA) was positioned 10 mm above the tube junction. Sonovue microbubble contrast agents (Bracco) were diluted in 2500 times and stirred by a magnetic stirrer to ensure the consistent microbubble concentration. The suspension was continuously injected into the inlet of the Y-shaped junction via syringe pump at 10 mL/h. Ultrasound RF data were acquired at 500 Hz frame rate for 40 s for further multi-stage SVD processing.

### 2.2. *In Vivo* Rat Brain Imaging

All animal experiments were performed in compliance with the European Community Council Directive of September 22, 2010 (2010/63/EU), approved by the local ethics committee (Comité d’éthique en matière d’expérimentation animale no. 59, “Paris Centre et Sud”), and conducted in accordance with the ARRIVE guidelines. In accordance with the 3Rs principles (replacement, reduction, and refinement), the number of animals was kept to the minimum necessary; accordingly, the animal used in this study was not euthanized for the purpose of the experiment but was included as part of an ongoing project (Project #2020-16). The experiment presented here was conducted prior to the scheduled euthanasia. Experiments were performed on a single Sprague–Dawley rat (Janvier Labs), weighing 300 g at the beginning of the experiments. The animal arrived at the laboratory one week prior to the experiment for acclimatization to the housing conditions and was maintained under controlled conditions (22 ± 1 °C, 60 ± 10% relative humidity, 12 h light/dark cycle, food and water ad libitum).Under deep anesthesia with intraperitoneal (i.p.) bolus of medetomidine (Domitor, 0.4 mg.kg^−1^) and ketamine (Imalgène, 40 mg.kg^−1^), a catheter filled with saline was inserted in the jugular vein of the rat before positioning the animal on the stereotaxic frame. The depth of anaesthesia was monitored throughout the imaging session from the heart (using ECG electrodes placed subcutaneously, AD Instruments & Labchart) and breathing rates (Spirometer, AD instruments, Labchart). The temperature of the animal was also monitored during the procedures using a rectal probe connected to a heating pad set at 37 °C (Physitemp, Clifton, USA). Eyes of mice were protected using a protective gel (Ocry-gel, TVM, UK). A sagittal skin incision was performed across the posterior part of the head to expose the skull. We excised the parietal and frontal flaps by drilling and gently moving the bone away from the dura mater. The opening exposed the brain from Bregma 0 mm to Bregma −8.0 mm. 1mL of saline, followed by 2mL of echographic were positioned on the rat brain. For ULM acquisitions, 0.15 mL of Sonovue microbubbles was injected via the jugular catheter, followed by 0.2 mL saline lush. Ultrasound RF data were acquired at 500 Hz for 10 seconds during the bolus injection.

### 2.3. Multi-Stage SVD Processing

For image acquisition for 2D imaging, ultrasound sequences were implemented and executed on a research ultrasound system (Verasonics, USA) driving a linear ultrasound transducer (128 elements, 15.625 MHz central frequency and a 67% bandwidth at −6 dB). All acquisition scripts and processing codes were developed in Matlab (Matworks, USA). For the HAM-SVD pulse sequence, single-cycle plane waves were transmitted at a frequency of 10 MHz at a frame rate of 500 Hz (A detailed calculation of theoretical frame rate analysis can be found in Supplementary material 1). Each transmit burst consisted of 5 plane-wave angles (−5°, −2.5°, 0°, +2.5°, +5°). Then the pulses were repeated for 4 different duty cycles (DC1 = 0.55, DC2 = 0.70, DC3 = 0.85, DC4 = 1.00) and 2 opposite polarities (+1 and −1). By varying the duty cycle, the transmit amplitude is modulated, enabling the acquisition of nonlinear responses at different pressure levels. Therefore, for each transmission, it consists of 40 plane waves (2 polarities × 5 angles × 4 DC cycles). The corresponding pulse repetition frequency (PRF) used in this study is 20,000 Hz. The ultrasound raw radio frequency (RF) data was acquired and stored for later image beamforming. For image beamforming, the RF data were offline beamformed using a delay-and-sum beamformer on GPU, utilizing a resolution grid with a spacing of 0.5 λ, resulting in the generation of IQ data for each angle and pulse polarity at the corresponding duty cycles. The resulting angle compounded IQ data for 4 duty cycles and 2 polarities were saved and input for later multi-stage SVD processing.

For image acquisition for 3D imaging, ultrasound sequences were implemented and executed on a research ultrasound system (Verasonics, USA) driving a row-column addressed (RCA) array (160 elements, 15.625 MHz central frequency and a 80% bandwidth at −6 dB). All acquisition scripts and processing codes were developed in Matlab (Mathworks, USA). In the HAM-SVD pulse sequence, 3 half-cycle plane waves were transmitted at a transmission of 10 MHz, with 10 angles for rows and columns respectively, with an angle range from −2.25° to 2.25°. Then the pulses were repeated for 4 different duty cycles (DC=0.55, 0.70, 0.85, and 1.00). The duty cycle is defined as the ratio of the pulse duration to the pulse repetition period, representing the fraction of time the transducer is actively transmitting during each cycle. Higher duty cycles result in greater energy transfer, which increases the effective average intensity of the transmitted wave. DC1-DC4 correspond to four different duty cycles, each resulting in a distinct transmit amplitude. By varying the duty cycle, the transmit amplitude is modulated, enabling the acquisition of nonlinear responses at different pressure levels. After the acquisition of ultrasound raw data, the radio frequency data were offline beamformed using a delay-and-sum beamformer on a GPU, utilizing a resolution grid with a spacing of lambda/2, resulting in the generation of IQ data.

For the first-stage spatial-angular SVD beamformer as described in details in [9], all the IQ dataset was reshaped into Nz, Nx, Nt, Np, Na, then the spatial-angular SVD was performed across all 5 angular frames for each Nt and Np. Only the first angular singular vector was retained to extract the most coherent signals in the angular domain.

For the second-stage spatial-pressure SVD processing, 8 frames at 4 duty cycles and 2 polarities were reshaped into a Casorati matrix for singular value decomposition processing. After the decomposition, the spatial singular vectors, U, pressure singular vectors, V, and the diagonal matrix, S, were obtained for SVD threshold selection. Spatial similarity matrix based on the spatial singular vectors can be used to observe the spatial coherence between each spatial singular vector [19]. During HAM-SVD, the singular vectors 1, 2, and 3 were separately selected to generate 3 individual datasets along the temporal domain. The HAM-SVD imaging datasets were stored and input for later spatiotemporal SVD filtering.

For the third-stage spatiotemporal SVD filtering, the first singular mode was discarded to avoid static tissue echoes, and all the other modes were combined to retrieve microbubble signals. After that, 1-lag autocorrelation and a rolling-basis temporal averaging across three frames were applied to further enhance the microbubble signals and de-noise [20]. Finally, the filtered datasets were summed up in the temporal direction to generate the final SVD-filtered Doppler images.

### 2.4. Multi-Modal SVD Analysis

To better interpret the outcomes of the SVD processing, we employed three complementary analyses: (1) the spatial similarity matrix (SSM) derived from the spatial singular vectors, *U;* (2) the logarithmic representation of the singular values in *S;* and (3) the Fourier transform of the temporal singular vectors, *V*. Together, these analyses provide a more comprehensive and intuitive visualization of the spatial and temporal characteristics of the decomposed modes.

Regarding the spatial singular vectors, *U*, spatial similarity matrix (SSM) was performed by calculating the correlation between all pairs of absolute values of the spatial singular vectors after normalization. Tissue and contrast subspaces can be observed from the SSM. A high-correlation block (top-left square) normally corresponds to tissue-dominated singular vectors, as tissue exhibits strong spatial coherence. A lower-correlation block represents contrast-dominated singular vectors, since contrast signal introduces spatial-pressure decorrelation, reducing pairwise correlations. The off-diagonal regions demonstrate the weak correlations (darker areas), which reflect dissimilarity between tissue and contrast subspace.

Regarding the diagonal matrix, *S*, we derived the energy spectrum by first normalizing the singular values (the diagonal of S) with respect to their maximum value and then converting them to a dB representation using 20 log10(.). This step highlights the dynamic range of the singular components.

Regarding the temporal singular vectors, *V*, Fourier transform (FFT) of temporal singular vectors was performed to evaluate the change in frequency at each temporal singular vectors. The Fourier transform of temporal singular vectors converts the amplitude domain signals into a frequency domain, revealing the spectral characteristics of the underlying components (tissue, microbubble signals, or noise). Each temporal singular vector represents the temporal evolution of a specific mode. Its Fourier transform shows the dominant frequencies associated with tissue motion (low-frequency), slowly-moving microbubbles (low-mid band), fast-moving microbubbles (mid-high frequency), noise (high-frequency), and also the energy distribution across frequencies for each mode.

### 2.5. Ultrasound Localization Microscopy

SVD-filtered images were interpolated (Lanczos interpolation kernel) down to (probe spatial λ/10 x λ/10). Microbubbles were detected as the brightest local maxima with high correlation (> 0.5) with a typical point spread function (harmonic imaging response of an isolated microbubble, modelled as a Gaussian spot of axial and lateral dimension of lambda). Sub-pixel maxima localization was then performed using a fast local second-order polynomial fit. The resulting coordinates were rounded to the chosen pixel size (here 15 × 15µm). Tracking of the maxima positions was performed using a classical particle tracking algorithm (simpletracker.m available on Mathworks ©Jean-Yves Tinevez, wrapping Matlab munkres algorithm implementation of ©Yi Cao 2009), with no gap filling and maximal linking distance corresponding to a 100mm/s maximum speed. Only tracks with MB detected in at least 30 successive ultrafast frames were selected. The successive positions gathered in one track were used to compute the interframe bubble velocity vector components (along probe x-axis and depth z-axis) and absolute velocity magnitude. We added a linear spatial interpolation on each track to count one MB detection in every pixel on the MB path. Maps of MB Count were computed by counting all the MBs detected in one pixel during the acquisition time; velocity maps were computed as their mean velocity. During the processing to create ULM images, microbubbles were detected as brightest local maxima with a good correlation with the PSF on filtered images. The value of this maximum was extracted on the filtered images and associated with each localized MB detection. This value is referred as the backscattering amplitude. Then, instead of attributing to each ULM map’s pixel the number of MB detected in this pixel during the whole acquisition, we attribute to each pixel the mean value of the backscattering amplitude of at the MBs detected in that particular pixel. The grid chosen is the same as the chosen one for the MB Count and Velocity maps.

### 2.6. Statistical Analysis

Statistical comparisons between datasets processed with conventional spatiotemporal SVD (C-SVD) and the proposed multi-stage SVD (MS-SVD) were performed using the Wilcoxon rank sum test (also known as the Mann–Whitney U test). This nonparametric test was chosen because the distributions of the evaluated metrics—microbubble flow speed, normalized correlation coefficient, backscattering amplitude — did not consistently satisfy the normality assumption required for parametric tests. For each comparison, the null hypothesis was that the two independent samples originated from distributions with equal medians. Two-tailed tests were applied, and p values less than 0.05 were considered statistically significant.

## 3. Results

### 3.1. *In vitro* 2D wall-less flow phantom imaging

Figure 3 summarizes the performance of MS-SVD compared with conventional spatiotemporal SVD in the Y-shaped wall-less flow phantom. A diluted SonoVue microbubble suspension was circulated at a controlled flow rate of 10 mL/h (theoretical velocity estimates in Supplementary Material 2), and RF data were acquired over 40 s at 500 Hz. Power-Doppler and ULM maps reconstructed from conventional SVD show visible background clutter and incomplete filling of the smaller branches, whereas MS-SVD produces much cleaner backgrounds and more continuous delineation of all three channels (Fig. 3a).

**Figure 1.**
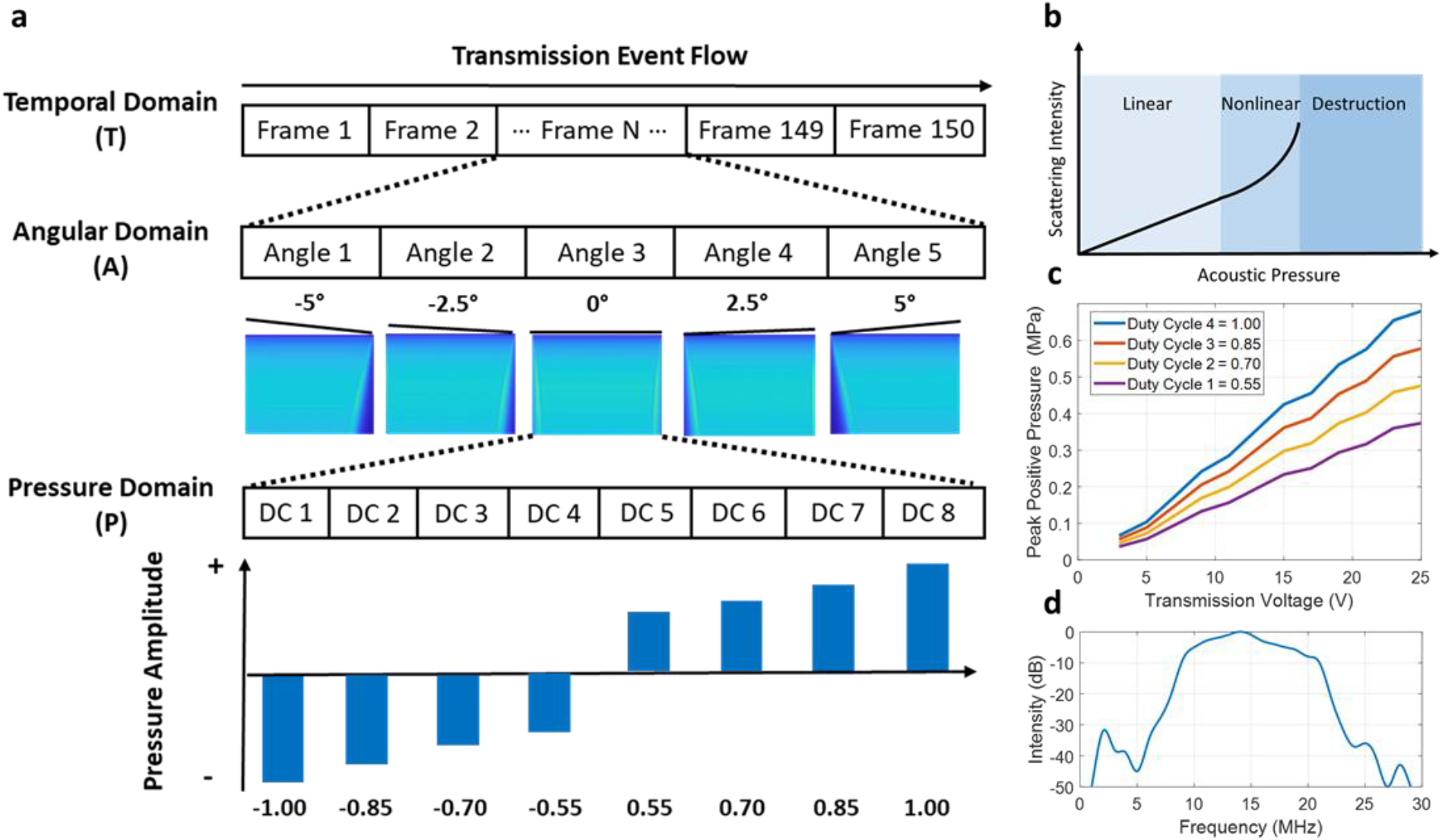
(a) MS-SVD transmission scheme: two half-cycle plane-wave was transmitted at 5 angles, for each angle, pulses were transmitted at 4 different duty cycles and 2 opposite polarities; (b) Schematic diagram of scattering intensity of microbubble as a function of the acoustic pressure; (c) Acoustic calibration of the pulses transmitted for MS-SVD with varying duty cycles used in this study; (d) Bandwidth of the transducer used in this study.

**Figure 2.**
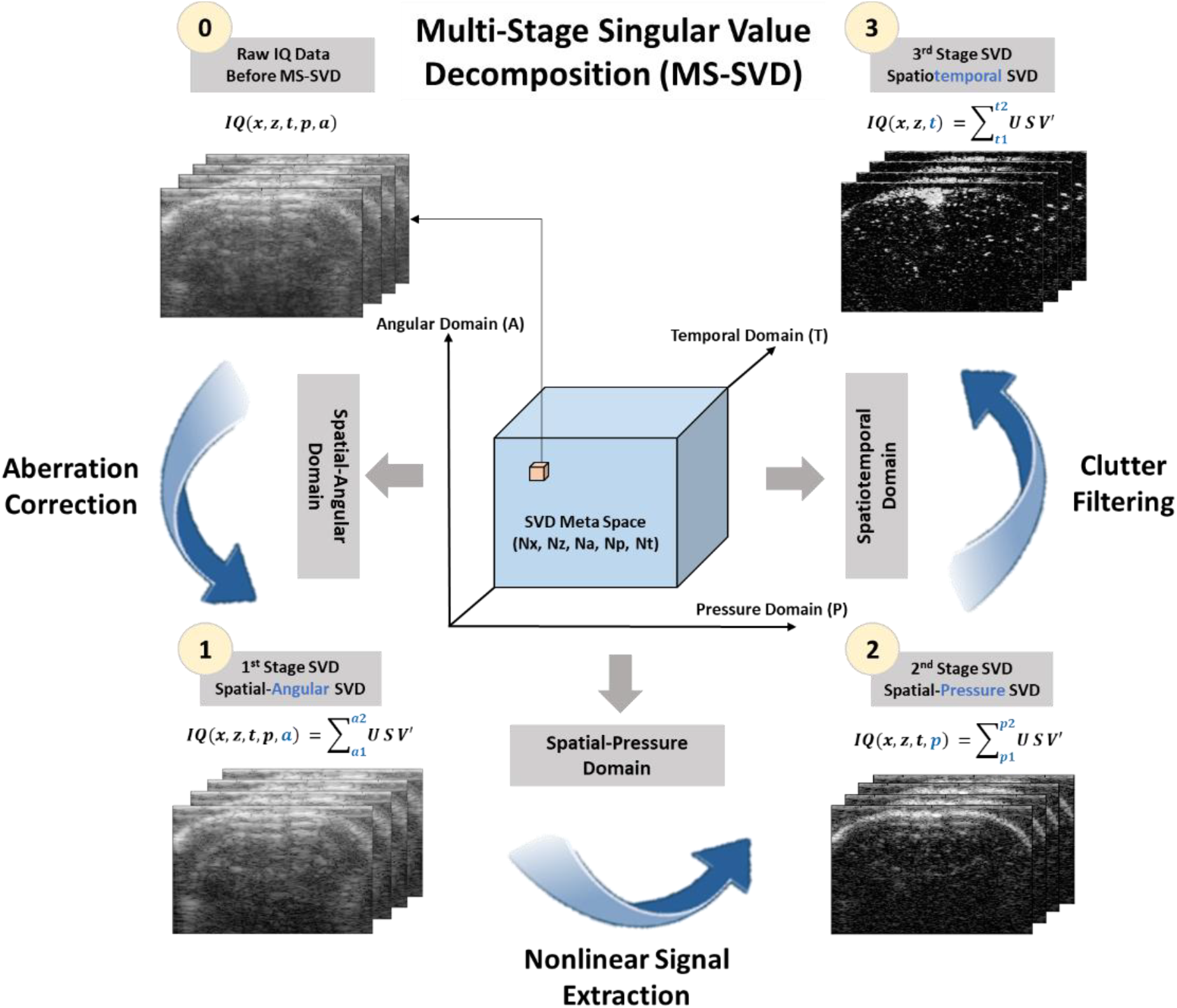
MS-SVD processing scheme: ultrasound data was acquired along pressure, angular, and temporal domains respectively, in order to generate a SVD meta space. Each voxel in this SVD meta space represent a single IQ image (Nx x Nz) at a certain angle (a), pressure level (p), and time point (t). First, the 1^st^ stage spatial-angular SVD was performed to extract the angular coherent signals; then the 2^nd^ stage spatial-pressure SVD was performed to extract the nonlinear contrast signals; finally, the 3^rd^ stage spatiotemporal SVD was performed to filter out the remaining tissue clutter signals, to generate final contrast imaging dataset for ultrasound localization microscopy.

**Figure 3.**
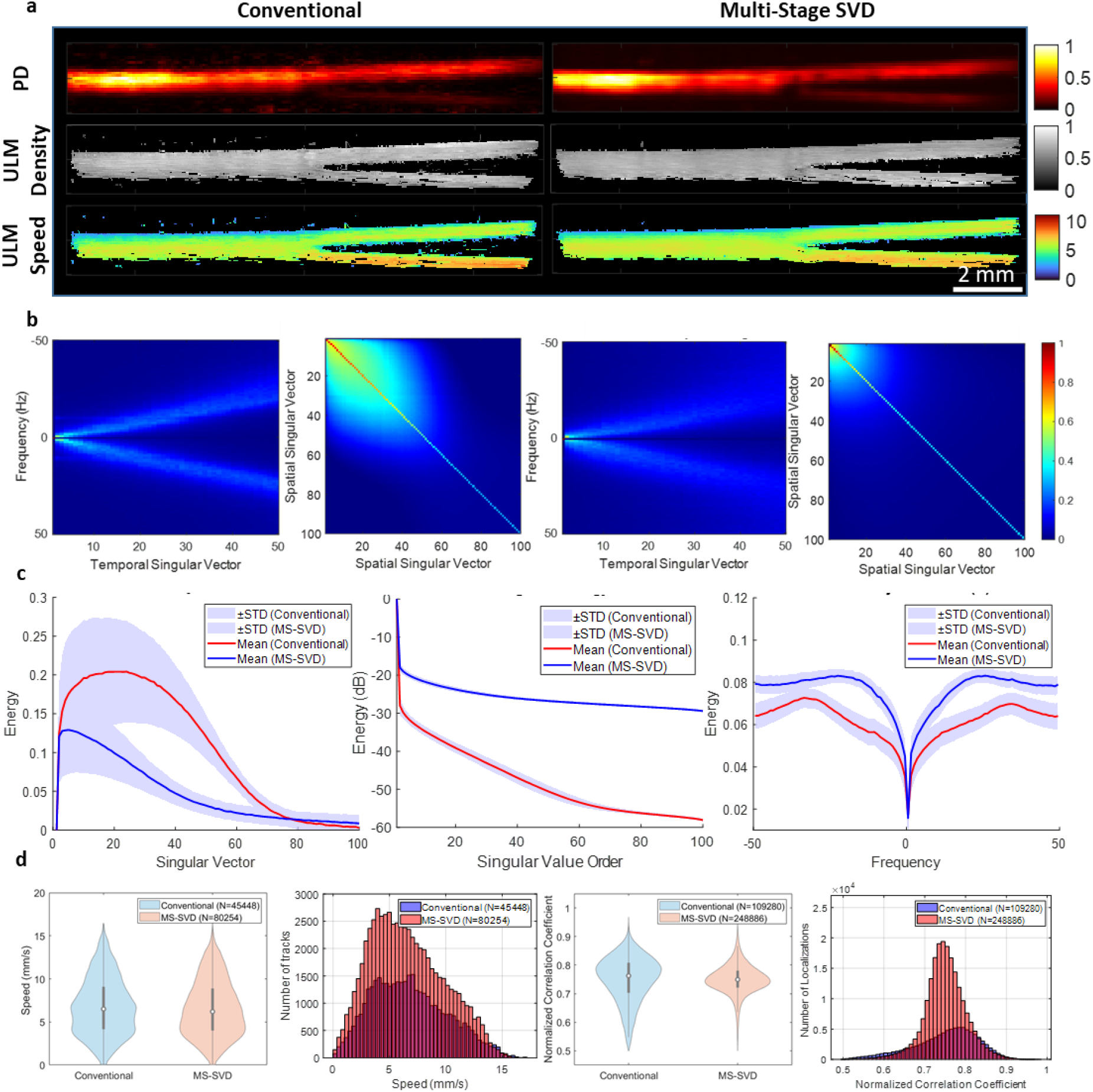
Comparison between conventional spatiotemporal SVD and MS-SVD for in vitro Y-shaped wall-less flow phantom. (a) Power-Doppler images, ULM localization density maps, and ULM speed maps obtained with conventional SVD and MS-SVD. (b) Fourier transform of the temporal singular vectors, V, and spatial similarity matrices based on the spatial singular vectors, U for both methods. (c) Axial projections of the spatial similarity matrices, logarithmic singular-value spectra (diagonal of S), and axial projections of the Fourier spectra of the temporal singular vectors. (d) Comparison of microbubble flow speed distributions and normalized correlation coefficient distributions between conventional SVD and MS-SVD in ULM for the Y-shaped phantom.

The multi-modal SVD analysis confirms that MS-SVD more efficiently concentrates coherent vessel signals into a limited set of modes. The Fourier spectra of the temporal singular vectors reveal that MS-SVD preserves more energy in the low-frequency band associated with slow bubble flow, while conventional SVD exhibits increased high-frequency content characteristic of residual clutter and noise (Fig. 3b). The spatial similarity matrices and their axial projections show that MS-SVD yields clear block-diagonal clusters, indicating well-separated tissue and contrast subspaces, whereas conventional SVD leads to more diffuse patterns (Fig. 3b–c). Consistently, the singular-value spectra decay more steeply for MS-SVD so that more than 80% of the energy is contained in roughly the first 40 modes, compared with about 60 modes for conventional SVD (Fig. 3c).

These improvements in clutter rejection translate directly into ULM performance. Super-resolved density and speed maps obtained after MS-SVD show denser, smoother filling of the Y-junction, with fewer gaps in low-contrast regions than the maps obtained from conventional SVD (Fig. 3a). Quantitatively, MS-SVD yields substantially more microbubble localizations in ULM than conventional SVD (248,886 vs. 109,280), and a higher number of successfully tracked trajectories (80,254 vs. 45,448), particularly for detections with normalized correlation coefficients between 0.7 and 0.8 (Fig. 3d). Overall, the phantom study indicates that MS-SVD enhances microbubble signal coherence across angles and pressures, sharpens single-bubble point spread functions (Supplementary Fig. 1), and provides higher-quality input for ULM than a single-stage spatiotemporal SVD.

### 3.2. *In vivo* 2D rat brain imaging

The proposed MS-SVD approach was next evaluated *in vivo* in a rat brain imaged through a cranial window after a SonoVue bolus injection (Fig. 4). Power-Doppler images averaged over 90 s (300 blocks) show that MS-SVD produces a cleaner and higher-contrast vascular map than conventional SVD (Fig. 4a). With MS-SVD, cortical vessels appear sharper and more continuous, and the parenchymal background is strongly suppressed, revealing small arterioles that are barely visible with conventional processing.

**Figure 4.**
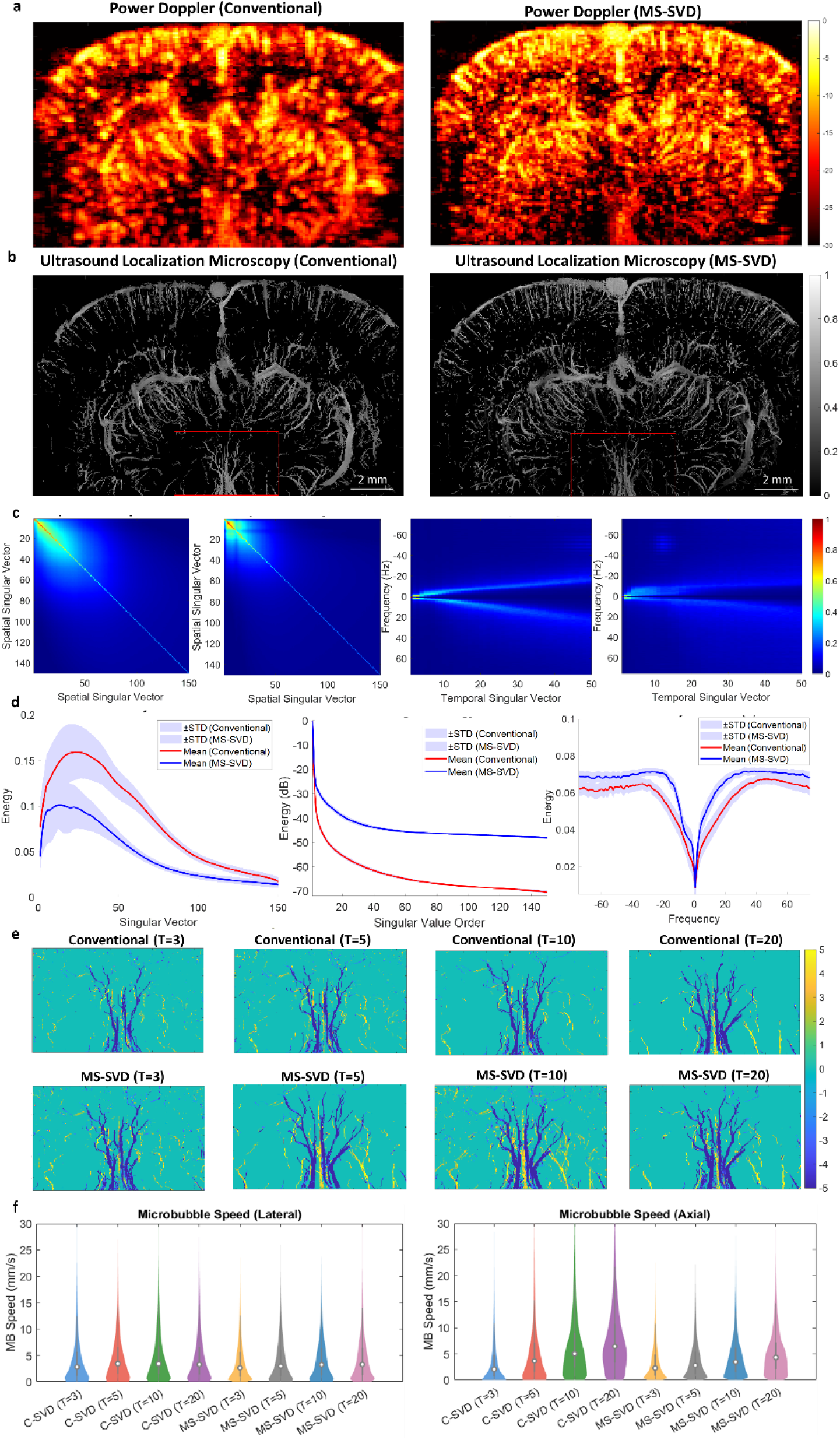
In vivo 2D rat brain imaging: comparison between conventional SVD and MS-SVD. (a) Power-Doppler images averaged over 90 s (300 blocks) obtained with conventional SVD and MS-SVD. (b) ULM super-resolution maps of the rat brain vasculature reconstructed from conventional SVD and MS-SVD filtered data. (c) Spatial similarity matrices derived from the spatial singular vectors, U, for both methods, singular-value energy curves from the diagonal of S, and Fourier spectra of the temporal singular vectors, V. (d) Histograms of microbubble flow speed estimated from ULM as a function of the starting SVD threshold. (e) Zoomed-in regions of the ULM reconstructions illustrating vessel sharpness and continuity for different starting SVD thresholds with conventional SVD and MS-SVD. (f) Quantification of microbubble flow speed components in the axial and lateral directions as a function of the starting SVD threshold T.

The SVD-derived metrics *in vivo* mirror the trends observed in the phantom. The spatial similarity matrix obtained from MS-SVD exhibits a pronounced block-diagonal structure, indicating that signals from individual vessels cluster into coherent singular-vector subspaces, whereas the conventional matrix remains more uniform and noisy (Fig. 4c). The singular-value spectrum for MS-SVD decays more rapidly than for conventional SVD, demonstrating that most of the microbubble flow energy is captured by only a few dominant modes (Fig. 4c). Likewise, the temporal spectra of the singular vectors show that MS-SVD concentrates energy at low frequencies (below approximately 5 Hz), consistent with slow cortical hemodynamics, while conventional SVD retains a high-frequency tail attributable to motion artifacts and residual noise (Fig. 4c–d and Supplementary Fig. 2).

ULM reconstructions from the *in vivo* data further highlight the advantages of MS-SVD. Super-resolution maps derived from MS-SVD exhibit a denser and finer microvascular network, with many small capillaries clearly delineated and vessel walls rendered with smoother contours than in the conventional SVD maps, which appear sparser and more pixelated (Fig. 4b,e). Zoomed-in regions confirm that MS-SVD processing yields continuous microbubble tracks and well-defined vessel cores, whereas conventional filtering leaves holes and spurious isolated detections (Fig. 4e). Quantitative analyses of ULM-derived metrics, including microbubble speed, show that MS-SVD recovers more microbubbles with higher coherence and signal strength and maintains more stable speed estimates across SVD thresholds (Supplementary Material 3 and related histograms and curves). Collectively, the 2D in vivo experiments demonstrate that MS-SVD substantially improves both Doppler-based and super-resolution readouts compared with a conventional spatiotemporal SVD filter.

### 3.3. *In Vivo* 3D Rat Brain Imaging

To assess the impact of MS-SVD in volumetric imaging, 3D acquisitions of the rat brain were performed and reconstructed using either conventional linear processing or the proposed MS-SVD pipeline (Fig. 5). Maximum-intensity power-Doppler projections show that MS-SVD yields a much clearer and more homogeneous visualization of the 3D vascular tree, with stronger contrast between vessels and surrounding tissue and improved continuity of deep cortical and subcortical branches compared with linear imaging (Fig. 5a). In particular, MS-SVD suppresses diffuse background speckle and unveils small vessels that are obscured or fragmented in the conventional reconstruction. The associated SVD-domain metrics again support more efficient clutter rejection with MS-SVD in 3D. The spatial similarity matrices derived from the spatial singular vectors show well-defined tissue- and contrast-dominated blocks for MS-SVD, whereas linear processing produces more mixed and less structured patterns (Fig. 5b). The singular-value energy curves exhibit a steeper decay for MS-SVD, indicating that most flow information is contained in the first few modes, and the Fourier spectra of the temporal singular vectors demonstrate that MS-SVD preferentially retains low-frequency content linked to slow microvascular flow while attenuating high-frequency clutter (Fig. 5b). ULM reconstructions performed on the 3D datasets highlight how MS-SVD benefits super-resolution volumetric imaging, especially for the transverse plane. Three-dimensional maximum-intensity projection maps reveal a denser microvascular network and finer capillary details in the MS-SVD-processed data than in the linearly processed volumes (Fig. 5c). Zoomed-in regions show that MS-SVD sharpens vessel boundaries and resolves closely spaced microvessels that remain blurred or merged under linear processing (Fig. 5d). Transverse slices through the volume confirm that MS-SVD-based ULM (nonlinear) provides vascular patterns that more closely match optical reference images than linear ULM, and fusion of linear and nonlinear reconstructions emphasizes complementary information (Fig. 5e). Finally, the distribution of microbubble speeds indicates that MS-SVD enhances sensitivity to slow-flow components while still capturing the full range of physiologically relevant velocities (Fig. 5f). Together, these 3D results demonstrate that MS-SVD extends its advantages to volumetric CEUS and ULM, enabling high-contrast, high-resolution mapping of rat cerebral microvasculature throughout the imaging volume.

**Figure 5.**
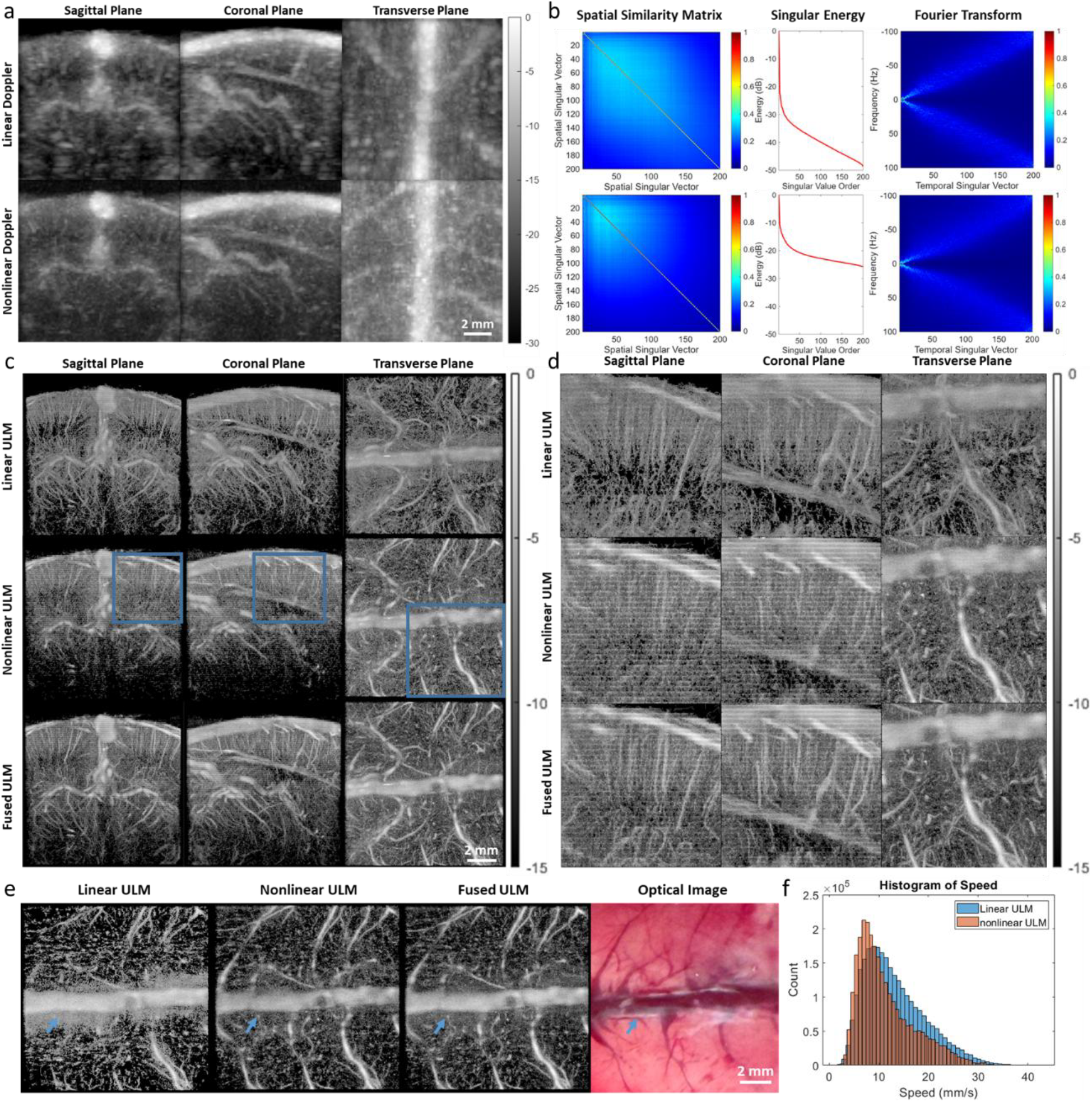
In vivo 3D rat brain imaging with conventional linear processing and MS-SVD (nonlinear) processing. (a) 3D power-Doppler maximum-intensity projections obtained with linear and MS-SVD processing. (b) Spatial similarity matrices from the spatial singular vectors, U, singular-value energy curves from S, and Fourier spectra of the temporal singular vectors V for linear and MS-SVD processing. (c) 3D ULM maximum-intensity projection maps of the cerebral vasculature reconstructed from linear and MS-SVD filtered data. (d) Zoomed-in views of selected regions (blue boxes in (c)) highlighting differences in vessel sharpness and capillary detail. (e) Transverse slices (3–4 mm depth) comparing linear ULM, nonlinear (MS-SVD) ULM, fused ULM images, and corresponding optical images. (f) Distributions of microbubble speeds measured with linear and nonlinear ULM reconstructions.

## 4. Discussion

We have introduced and validated a cascaded SVD imaging pipeline that significantly improves ultrafast CEUS of microbubbles. The central innovation is the multi-stage, multi-dimensional SVD strategy applied to data acquired with a transmit scheme (multiple angles, amplitudes, and time points). Each stage contributes to isolating microbubble signals from tissue:

- Stage 1 – Spatial-Angular SVD: By combining all compounding angles into a single angular mode, we exploit coherence across angles to ensure that the subsequent processing works on data where the signals are already coherently aligned in the angular domain.
- Stage 2 – Spatial-Pressure SVD (HAM-SVD): We reshape the data across the four pressure levels and two polarities and perform SVD in this domain. Microbubbles and tissue respond differently to amplitude modulation: for example, bubbles produce strong nonlinear second-harmonic echoes while tissue remains mostly linear.
- Stage 3 – Spatiotemporal SVD: Finally, we apply SVD in slow time on each of the separated datasets. Here the goal is to remove any remaining static or noisy signals, isolating true moving microbubbles. By using a relatively low rank (discarding only the top two singular values), we preserve slowly moving bubbles – something conventional temporal filtering often loses. This stage boosts the signal-to-noise ratio by discarding incoherent noise over time.

The combined effect of these stages is a remarkable enhancement of microbubble signal extraction. Our in vitro and in vivo experiments show that MS-SVD consistently suppresses tissue background much more strongly than single-stage methods. For example, in the Y-phantom the MS-SVD Doppler CNR was visibly higher and the PSFs were sharper (Fig. 3, Suppl. Figs. 1–2). In the rat brain, MS-SVD yielded clearer vessel contrast and retained more low-frequency energy associated with flow (Fig. 4c & 5b). These observations confirm that each SVD domain (angular, pressure, temporal) adds a layer of filtering that, together, yields much better clutter removal and microbubble preservation.

In terms of microbubble flowing speed analysis, as can be seen from the Fourier transform of the temporal singular vectors for in vitro (Fig 3b) and in vivo (Fig 4c) experiments that, MS-SVD demonstrated stronger energy accumulated in low frequency region compared to the conventional SVD. This finding also corresponds to the microbubble flowing speed derived from ULM (shown in Fig 4f & 6d and supplementary material 3). For the in vitro experiments, MS-SVD demonstrated a 3% lower microbubble flowing speed compared to the conventional SVD method, whereas for the in vivo study, MS-SVD showed 6%-29% lower microbubble flowing speed compared to the conventional SVD at different starting SVD threshold. This indicates MS-SVD captures more of the slowly-moving microbubbles. Additionally, we found out that, no matter for MS-SVD or conventional SVD methods, as the SVD starting threshold increases, the microbubble flowing speed will increases accordingly in ULM. More specifically, we looked at the speed components in the lateral and axial directions separately for in vivo rat brain imaging, as shown in Fig 4f. It indicated that, the speed variation at different SVD thresholding originated from the microbubble flowing speed in the axial direction. The possible reasons could be due to the beam-flow angle, which leads to a more sensitive flow estimation in the axial direction. For the quantification of normalized correlation coefficients, although MS-SVD and conventional SVD methods demonstrated different distribution of normalized correlation coefficients in the histogram as shown in Fig 3d for the in vitro phantom experiment. They showed the same mean value of 0.75 as shown in the supplementary material 3. Importantly, the normalized correlation coefficient distribution of the MS-SVD is narrower and centered around high values while conventional SVD leads to widely spread distribution of normalized correlation coefficient.

In general, MS-SVD demonstrated a significantly smaller PSFs compared to conventional method. However, there are a sub-population of microbubble signals whose intensities are relatively low on spatial-pressure SVD filtered images. There are a few possible explanations for this. First, the bandwidth of the transducer is from 10.4 - 20.8 MHz at −6 dB. Since the transmission was set at 10 MHz in this study, the second harmonic signals should fall at around 20 MHz. Therefore, there may be some parts of second harmonic signals which cannot be completely received due to the bandwidth. Second, according to the simulation of the acoustic field in the elevational direction, as the increase of the duty cycles, it can be seen that, the thickness of the acoustic beam in the elevational direction changes. However, for the spatial-pressure HAM-SVD processing, it can only obtain the nonlinear harmonic signals which were covered by all transmissions in the elevational directions. Therefore, the imaging plane of using HAM-SVD sequence may be thinner in the elevational direction compared to the conventional B-mode imaging sequence.

Compared to prior approaches, MS-SVD also retains important practical advantages. Because it is based on plane-wave imaging, it preserves the ultrafast frame rate and large field-of-view of ultrafast CEUS. Unlike line-by-line methods (e.g. pAM/xAM), our protocol uses simple global duty-cycle modulation, avoiding the need for complex element-by-element coding. This simplicity comes with one trade-off: our multi-angle, multi-amplitude transmit sequence requires a programmable research scanner. Not all clinical machines allow rapid switching of amplitude and polarity in this way, which may limit immediate translation.

Another practical challenge is threshold selection in SVD filtering. The cutoff between signal and clutter is critical: too aggressive filtering would discard slow microbubbles, while too lenient filtering leaves noise. In this study, we chose to keep only the largest singular values (e.g. first two) in the final stage to avoid losing slow flows. Our spatial-pressure filtering similarly used the identified harmonic component(s) from a fixed singular index. In future work, data-driven or adaptive threshold methods could further optimize this trade-off for different scenarios.

Computational load is also a consideration: the 3-stage SVD pipeline is intensive, as it processes large 4D datasets (space × angles × pressures × time). We performed all SVDs offline on a GPU-accelerated system, but real-time imaging would require further acceleration. Techniques such as randomized SVD or dedicated hardware (FPGAs/GPUs) could reduce latency.

Despite these challenges, the benefits of MS-SVD are clear. The greatly improved microbubble sensitivity and SNR have significant implications for super-resolution ultrasound. By providing a cleaner, higher-contrast input to localization algorithms, MS-SVD can markedly enhance ULM outcomes. In our experiments, MS-SVD enabled detection of many more bubbles and resolved finer capillaries (Fig. 6). This means that in practice, MS-SVD can shorten acquisition time or improve resolution for ULM. More broadly, any CEUS application that relies on detecting weak nonlinear signals (such as molecular imaging or perfusion studies) can benefit from MS-SVD’s enhanced contrast specificity.

## Conclusion

We have developed a comprehensive multi-stage SVD framework that greatly advances ultrafast CEUS of microbubbles. By combining a specialized multi-parameter transmit sequence with a cascade of SVD filters across spatial, angular, pressure, and temporal domains, the method isolates the complete spectrum of microbubble echoes (fundamental and second-harmonic) from tissue clutter. Our phantom and in vivo results show that MS-SVD significantly outperforms conventional filtering: it achieves much stronger tissue suppression, higher contrast-to-noise ratio, and many more bubble detections. In practice, this yields denser super-resolution maps of microvasculature. Crucially, MS-SVD retains the high frame rate and wide field-of-view of ultrafast imaging while overcoming key limitations of prior clutter-filtering schemes.

Looking forward, MS-SVD opens the door to more sensitive and reliable microvascular imaging. The enhanced input it provides to ultrasound localization microscopy should translate into finer and faster super-resolution imaging. the present work establishes multi-stage SVD as a powerful new front-end for high-sensitivity CEUS and super-resolution ultrasound. Our findings represent a substantial step toward routine high-contrast visualization of slow microcirculation in research and clinical ultrasound.

## Acknowledgements

This work was supported by Inserm research accelerator (Inserm ART) in Biomedical Ultrasound and by the French national research agency (ANR) under ANR-21-CE19-0050 program (Project SonoGT). This research was also funded by the Region Ile de France - Convention DIM ELICIT Innovative Technologies for Life Science. This work was also partially funded by the AXA research fund (Project NeuroElastoFlow).

## Data Availability

Data supporting the findings of this study are available in the framework of an official collaboration between academic institutions. Correspondence and requests for data should be addressed to M.T.

## Supplementary Material 1

### Theoretical Frame Rate Analysis

For a 10mm depth acquisition and number of pressure amplitude N=4 (achieved by 4 different duty cycles), theoretically acquired along the 128 elements of a linear array transducer. One single plane-wave acquisition is then acquired in:

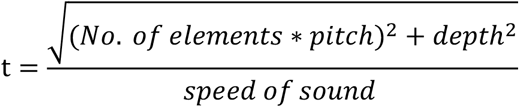

with number of elements = 128, pitch = 0.10 mm, depth = 10mm, and speed of sound = 1540 m/s

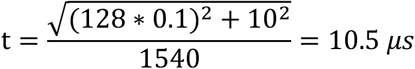

then, one image of the proposed HAM-SVD method (plane-wave transmissions across 11 angles at 4 different pressure amplitudes) and 2 different polarities can theoretically be acquired in:

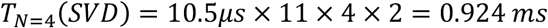

Which corresponds to an imaging frame rate of 1.08 kHz. (As it can be achieved by using a single voltage, therefore, there is no need to change TPC profile in the Verasonics system)

## Supplementary Material 2

### Theoretical Flow Velocity Analysis

For the Y-shaped junction tube used for the in vitro tube phantom experiments:

Since the inlet is a single tube with a diameter of 0.8 mm and the outlet has two branches, each of diameter of 0.4 mm. The total flow rate from the inlet was set at 10 mL/hour, theoretically evenly split between two outlets.

Then, the conversion of flow rate:

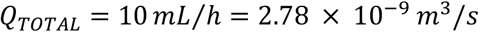

Calculation of the cross-sectional areas and the corresponding flow velocities:

For the inlet with the diameter of 0.8 mm:

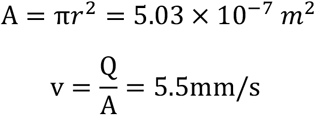

For each outlet with the diameter of 0.4 mm:

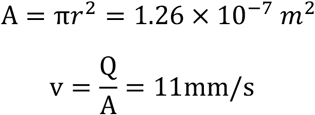

## Supplementary Material 3

**Supplementary table 1.** Comparison of mean and standard deviation of speed in mm/s between using C-SVD and MS-SVD for the in vivo rat brain imaging at different SVD starting thresholds T.

| Speed | T=3 | T=5 | T=10 | T=20 |
| --- | --- | --- | --- | --- |
| C-SVD | 6.19 ± 5.05 | 7.84 ± 5.43 | 9.06 ± 6.04 | 9.94 ± 6.23 |
| MS-SVD | 5.84 ± 4.38 | 6.43 ± 4.47 | 7.03 ± 4.58 | 7.67 ± 4.61 |
| Difference | 6.0 % | 21.9 % | 28.9% | 29.6 % |
| P-value | 0.001 | 0.001 | 0.001 | 0.001 |

**Supplementary table 2.** Comparison of mean and standard deviation of speed (in the lateral direction) in mm/s between using C-SVD and MS-SVD for the in vivo rat brain imaging at different SVD starting thresholds T.

| Speed (Lateral) | T=3 | T=5 | T=10 | T=20 |
| --- | --- | --- | --- | --- |
| C-SVD | 4.16 ± 4.25 | 4.68 ± 4.30 | 4.80 ± 4.50 | 4.67 ± 4.46 |
| MS-SVD | 3.89 ± 3.85 | 4.14 ± 3.92 | 4.36 ± 4.02 | 4.37 ± 4.02 |
| Difference | 6.9 % | 13.0 % | 10.1% | 6.9 % |
| P-value | 0.001 | 0.001 | 0.001 | 0.001 |

**Supplementary table 3.** Comparison of mean and standard deviation of speed (in the axial direction) in mm/s between using C-SVD and MS-SVD for the in vivo rat brain imaging at different SVD starting thresholds T.

| Speed (Axial) | T=3 | T=5 | T=10 | T=20 |
| --- | --- | --- | --- | --- |
| C-SVD | 3.48 ± 4.05 | 5.11 ± 4.99 | 6.55 ± 5.76 | 7.78 ± 5.96 |
| MS-SVD | 3.38 ± 3.49 | 3.86 ± 3.74 | 4.44 ± 4.02 | 5.21 ± 4.20 |
| Difference | 2.9 % | 32.4 % | 47.5% | 49.3 % |
| P-value | 0.001 | 0.001 | 0.001 | 0.001 |

**Supplementary table 4.**
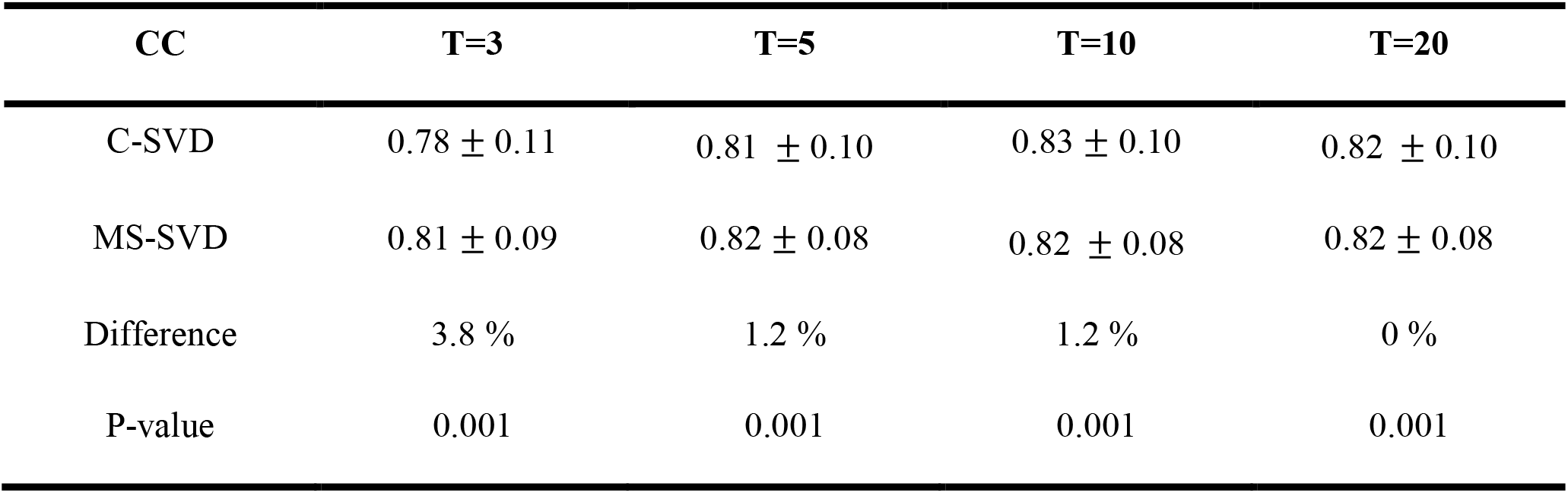
Comparison of mean and standard deviation of normalized correlation coefficient between using C-SVD and MS-SVD for the in vivo rat brain imaging at different SVD starting thresholds T.

**Supplementary table 5.** Comparison of mean and standard deviation of backscattering amplitude between using C-SVD and MS-SVD for the in vivo rat brain imaging at different SVD starting thresholds T.

| BSA | T=3 | T=5 | T=10 | T=20 |
| --- | --- | --- | --- | --- |
| C-SVD | $215 \pm 168$ | $149 \pm 133$ | $104 \pm 91$ | $100 \pm 82$ |
| MS-SVD | $205 \pm 198$ | $179 \pm 172$ | $145 \pm 136$ | $110 \pm 98$ |
| Difference | 2.9 % | 32.4 % | 47.5% | 49.3 % |
| P-value | 0.001 | 0.001 | 0.001 | 0.001 |

**Supplementary table 6.** Comparison of mean and standard deviation of microbubble flowing speed, normalized correlation coefficient, and backscattering amplitude between using C-SVD and MS-SVD for the in vitro wall-less flow phantom at a SVD starting threshold of T=2.

| In Vitro | Speed | NCC | BSA |
| --- | --- | --- | --- |
| C-SVD | $6.79 \pm 3.30$ | $0.75 \pm 0.08$ | $7.28 \pm 2.88$ |
| MS-SVD | $6.58 \pm 3.25$ | $0.75 \pm 0.05$ | $6.57 \pm 3.40$ |
| Difference | 3.2 % | 0 % | 10.8% |
| P-value | 0.001 | 0.94 | 0.001 |

## Supplementary Material 4

**Supplementary Figure 1.**
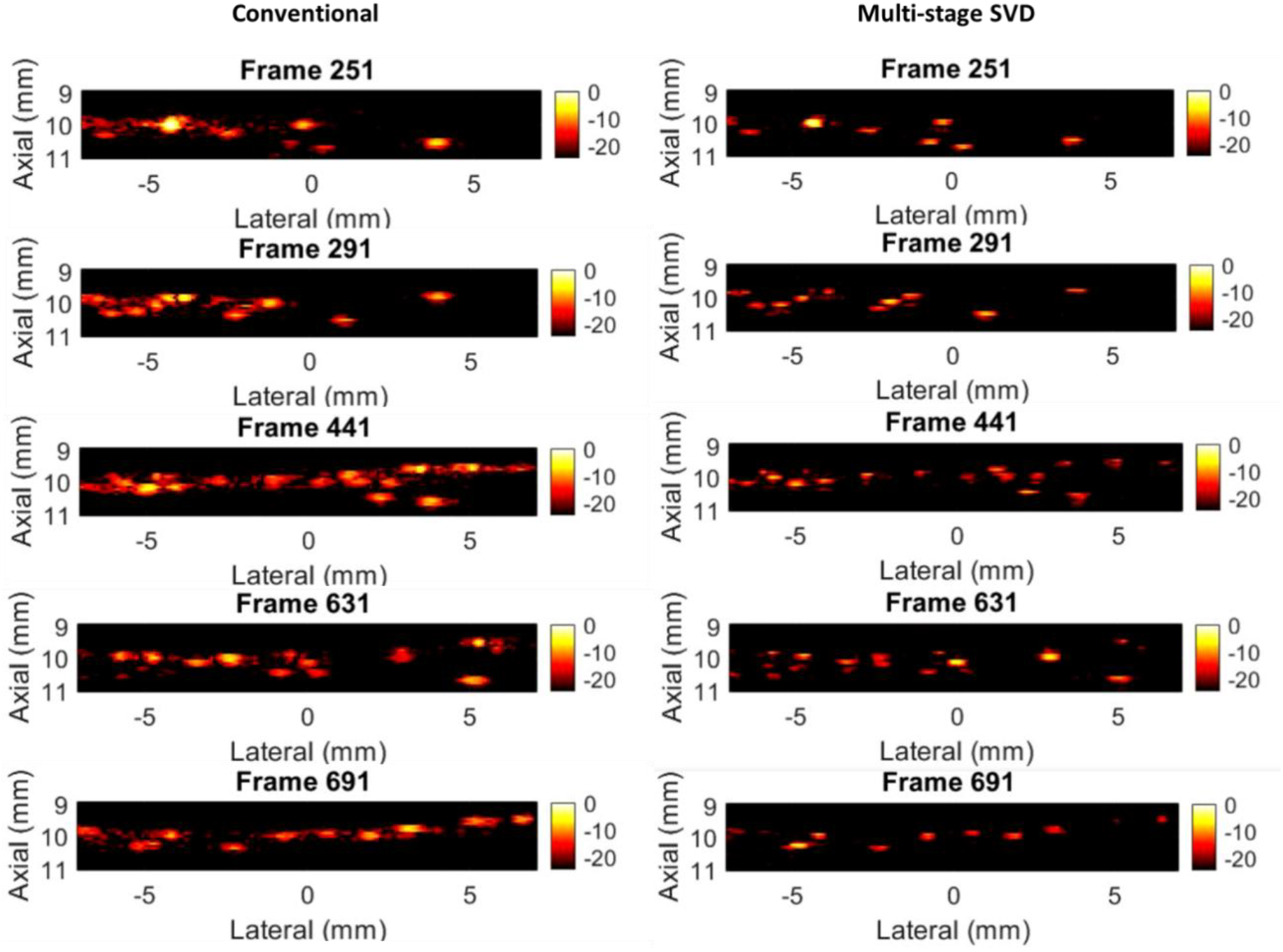
The comparison of single frames of conventional and Multi-stage SVD for Y-shaped tube phantom.

## Supplementary Material 5

**Supplementary Figure 2.**
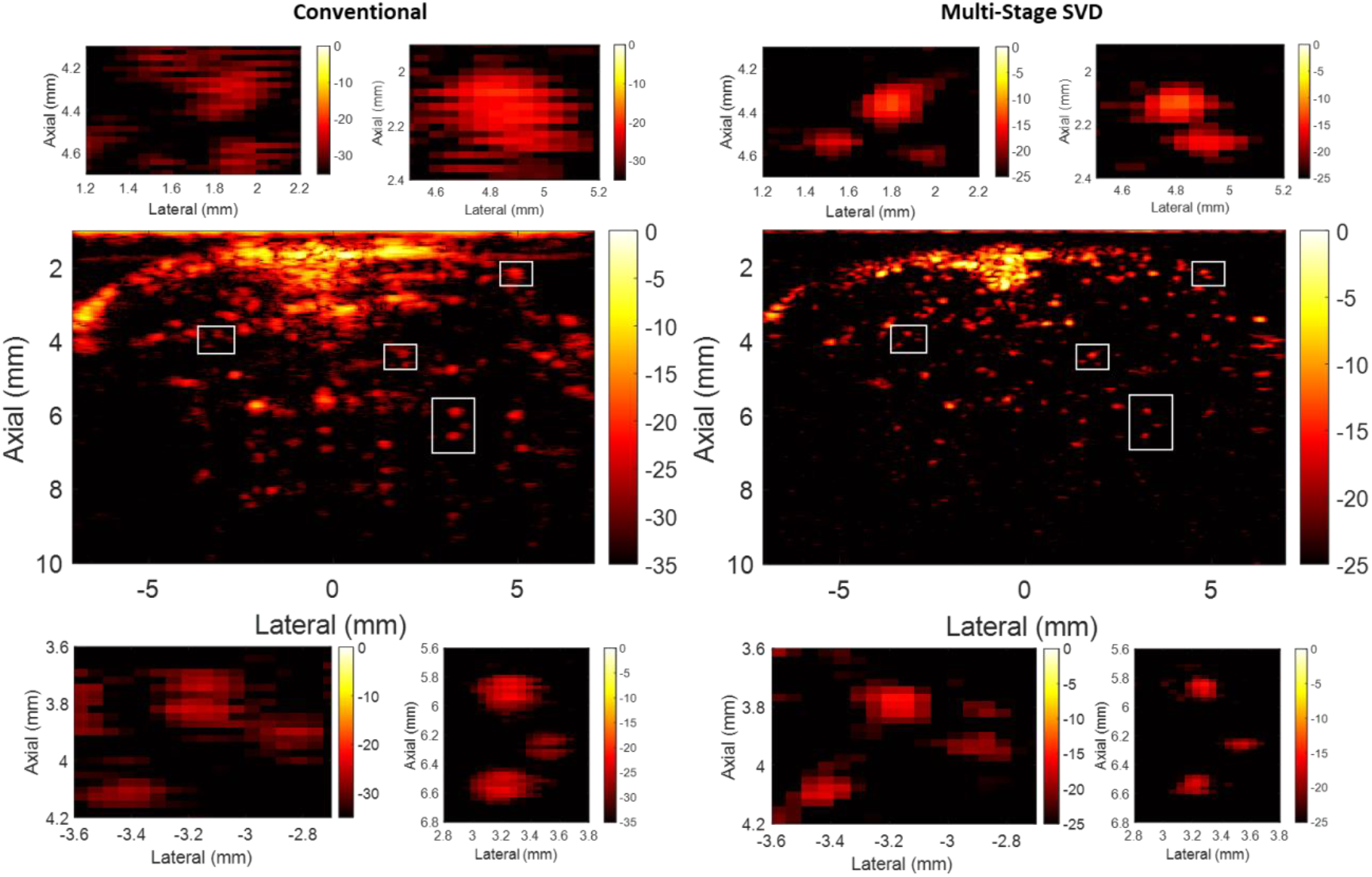
The comparison of single frame of conventional and Multi-stage SVD for the rat brain imaging.

## Supplementary Material 6

**Supplementary Figure 3.**
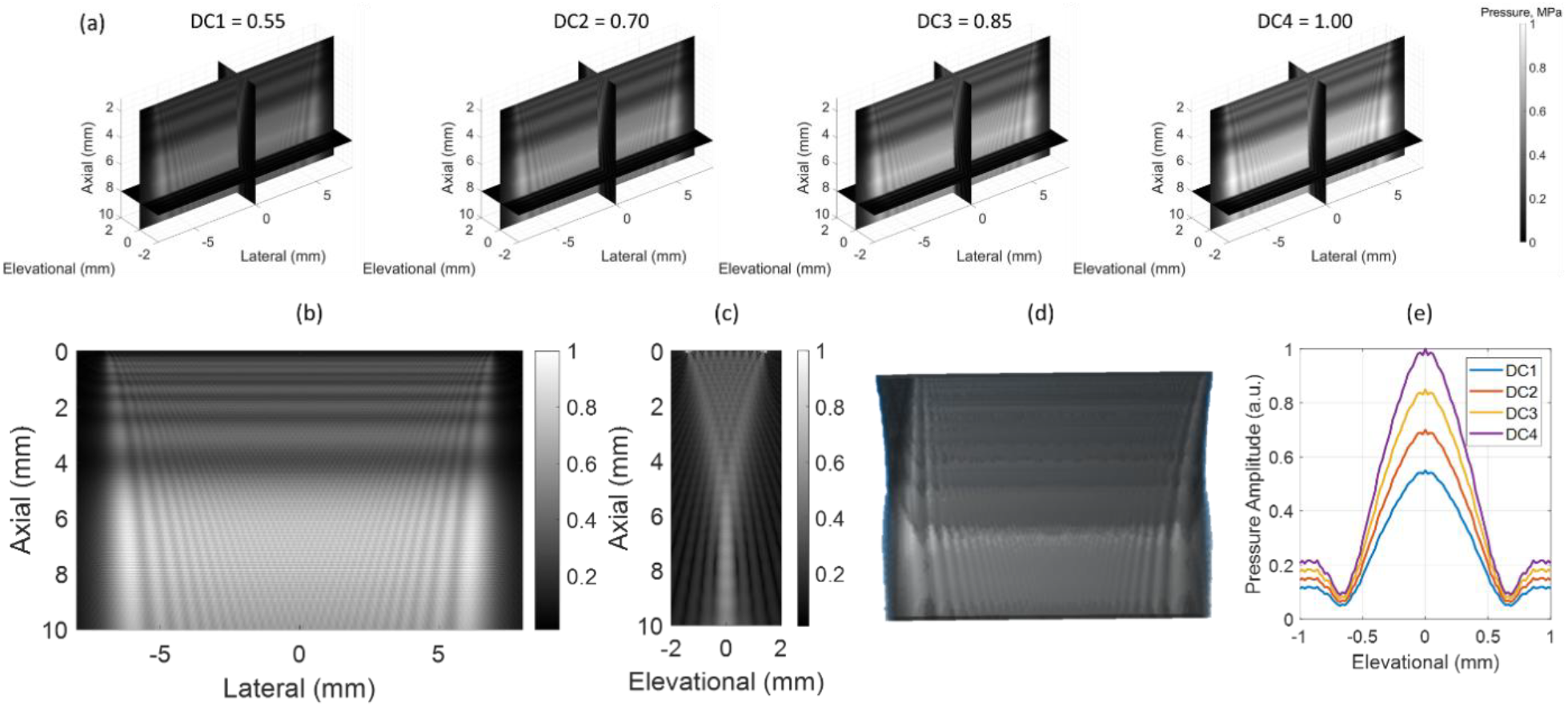
(a) Simulation of the acoustic pressure field across 4 different duty cycles. (b) Simulation of the acoustic pressure field in the axial-lateral direction (c) Simulation of the acoustic pressure field in the axial-elevational direction (d) volumetric visualization of the acoustic pressure field (e) Quantification of the acoustic pressure amplitude measured at the axial depth of 6 mm when transmitting 4 different duty cycles stated in this study.

## Supplementary Material 7

**Supplementary Figure 4.**
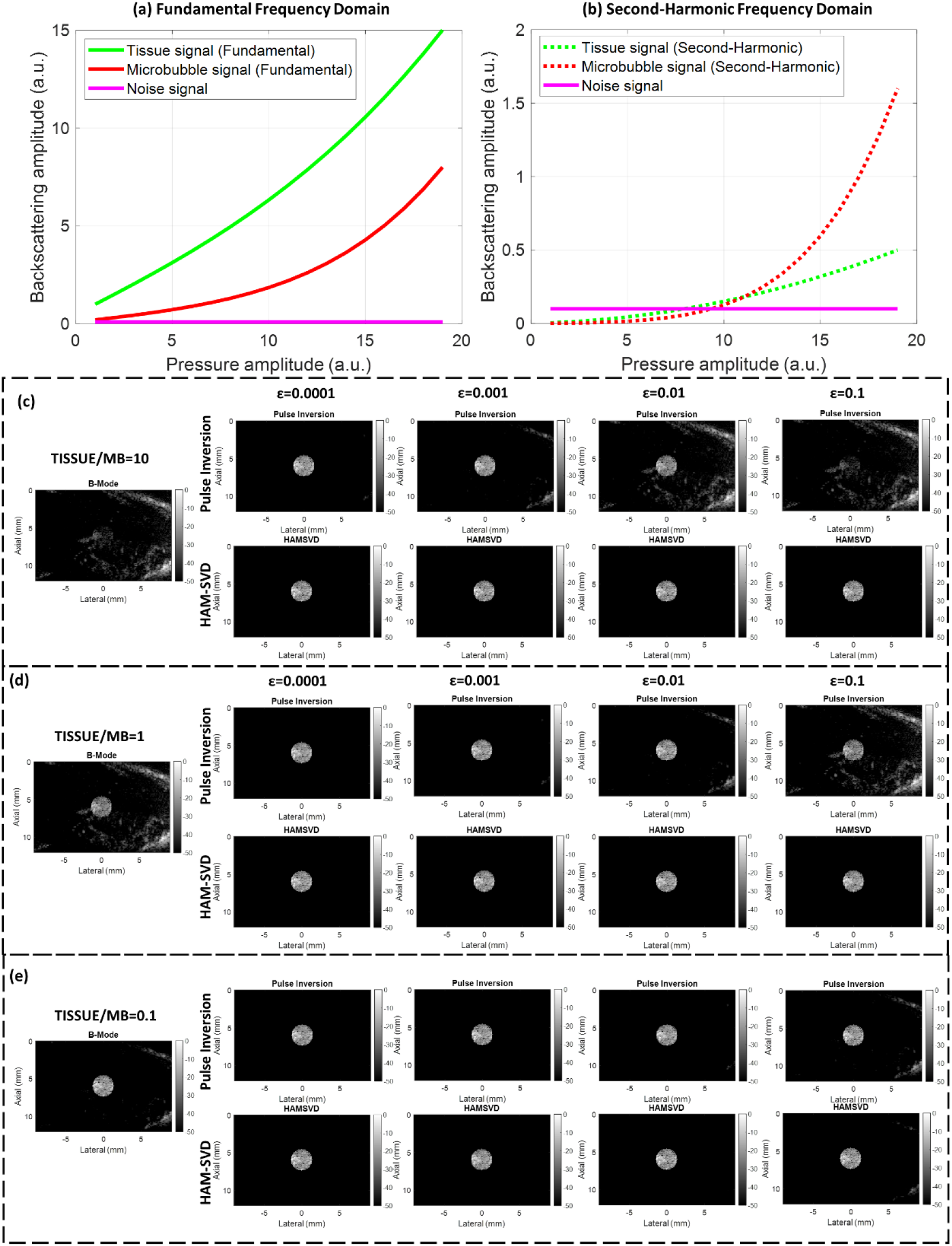
The relationship between backscattering amplitude of tissue and microbubble signals were simulated according to the changing pressure amplitude (a) in the fundamental frequency domain; (b) in the second-harmonic frequency domain; (c-e) Comparisons between pulse inversion and spatial-pressure SVD for different ε (ε represents the difference between positive and negative pulses) and ratios between the backscattering amplitudes of tissue and microbubbles (ratio=10, 1, and 0.1 respectively)

The relationship between backscattering amplitude of tissue and microbubble signals were simulated according to the changing pressure amplitude, in the fundamental frequency and second-harmonic frequency domains respectively, as illustrated in Supplementary Figure 5.

A tissue phantom with microbubble contrast was simulated using Matlab (Mathworks, USA), as depicted in Supplementary Figure 4. The final simulated in-phase quadrature (IQ) data, is basically comprised of three matrices describing the spatial distribution of tissue signal, S_T_, microbubble signals, S_MB_, and random noise signals, S_N_, respectively. The backscattering amplitudes of microbubbles, B_MB_, and tissue signals, B_T_, were simulated with respect to the pressure amplitudes, *p*. Thus, the IQ data can be mathematically expressed as function of spatial and pressure variables:

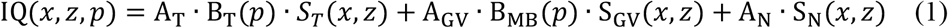

For the backscattering amplitude, both acoustic response in fundamental and harmonic responses were taken into account for both tissue and microbubble signals:

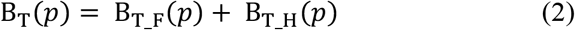

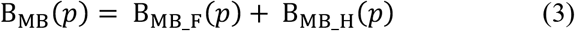

In fundamental frequency domain, tissue signal consists of strong linear signals and weakly nonlinear signals. For microbubble signals, it exhibits both the linear signals and also higher degrees of nonlinearity as the acoustic pressure increases.

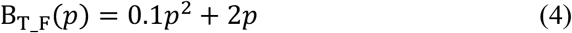

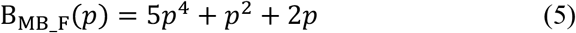

In the second-harmonic frequency domain, it is assumed that tissue signal does not have any linear signals and only weakly nonlinear harmonic signal left. For the microbubble signal, since it can generate the harmonic signals, so it will generate an even higher degree of nonlinearity as the acoustic pressure increases compared to the fundamental frequency domain.

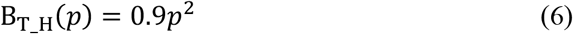

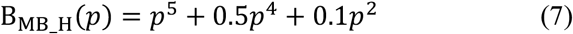

So if we put equation (4-7) into (1) and reshape the equation according to the power of p, we can have:

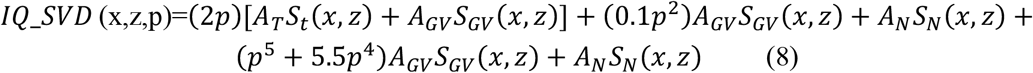

If we applied SVD on the equation (8), SVD can sort the signals in terms of different degrees of nonlinearities, so that we can have: Singular vector 1 = 2p (linear fundamental signal); Singular vector 2 = 0.1*p*^2^ (nonlinear fundamental signal); Singular vector 3 = *p*^5^ + 5.5*p*^4^ (nonlinear second-harmonic signal); Singular vector 4-8 = c (noise).

